# Sex-specific regulation of macrophage function determines outcome to acute bacterial infection

**DOI:** 10.64898/2026.08.28.747827

**Authors:** Rebecca L Belmonte, Julia Aleksandrowicz, Caroline Pumpe, Paulina Mika, Mary-Kate Corbally, David F Duneau, Jennifer C Regan

**Author notes:** Equal contribution (David F Duneau, Jennifer C Regan).

## Abstract

Sex differences in infection outcome are widespread across sexually reproducing animals. The immune mechanisms generating these differences remain incompletely understood, in part because it is challenging to decompose systemic effects from cell-intrinsic regulation in mammalian model systems and clinical data. Sex-biased infection outcomes are observed in taxa lacking adaptive immunity, suggesting that innate immune cells, such as macrophages, have the potential to drive dimorphisms. It is largely unknown whether macrophage-intrinsic sex identity is causal for infection susceptibility, or for sex differences in other, homeostatic functions. Here, we address this question using *Drosophila melanogaster*, an *in vivo* model of innate immunity where sex is established, and can be manipulated, cell-autonomously. We show that during infection by the bacterium *Staphylococcus aureus*, adult male flies succumb faster than females, with more rapid early bacterial proliferation. Hemocyte ablation reveals that survival is hemocyte-dependent in both sexes, and flow cytometry indicates a higher fraction of actively phagocytic hemocytes in females. Critically, genetically feminizing male hemocytes abolishes the dimorphism in survival and bacterial burden in *S. aureus* infection, bringing feminized males to the equivalent load and mortality risk as females. Transcriptomic analysis of whole carcasses shows strongly sex-biased responses during *S. aureus* infection, whereas hemocyte-specific RNA-sequencing reveals minimal sex differences in induced responses but substantial, sustained baseline transcriptomic divergence, including female-biased expression of bactericidal mechanisms linked to ROS generation and lysozyme production. Together, these findings demonstrate that the sex identity of innate immune cells is sufficient to shape infection outcome.

## Introduction

Sexual dimorphism in infection outcome is a pervasive feature of sexually reproducing taxa. Much of our understanding of this dimorphism derives from research on adaptive immunity in mammals, where sex hormones, such as estrogen and testosterone^1^, and karyotype, resulting X-linked gene dosage, contribute to distinct immune regulation in males and females^2^. Yet sex-biased infection outcomes are observed broadly across animal taxa, including those lacking adaptive immunity and sex steroid hormones ^3,4^, suggesting that innate immune cell function is regulated by sex. In mammals, cell autonomous sex differences in innate cell function are difficult to disentangle from the systemic effects of sex hormones and sexually dimorphic adaptive responses, which may obscure or amplify them. Macrophages exemplify this challenge: they show sex-biased differences in number, transcriptional state, and inflammatory function ^5–9^, yet whether these differences are themselves sufficient to determine infection outcome remains largely unknown. Resolving this is fundamental to explaining why males and females differ so consistently in their ability to combat infection, and could offer new insight into sex differences in innate cells both in health and disease states, including dysregulated inflammation.

Mammalian macrophages display sex differences in homeostatic functions, and responses to infection. In mice, peritoneal macrophages differ between sexes in their rate of replenishment and transcriptional state^5,7^. One consequence is sex-biased expression of the C-type lectin CD209b, which contributes to greater resistance to *Streptococcus pneumoniae* peritonitis in females^5^. Sex-specific regulation extends to macrophage survival and differentiation, with RELM-α required for these processes in females but not males^6^. Dimorphisms are observed beyond bacterial infection: female-biased outcomes following influenza virus infection are linked to differential innate immune responses including macrophage activation^2,10^. That sex-biased regulation of macrophages is at least partly cell-autonomous is supported by *in vitro* work in humans: monocyte-derived macrophages from male donors show higher *Leishmania infantum* burdens and a weaker type I interferon induction than those from female donors, even when cultured in hormone-stripped conditions^11^. However, such studies cannot control for host genetic or hormonal variation, and *in vivo* work cannot disentangle macrophage-intrinsic sex effects from circulating sex hormones and sexually dimorphic adaptive immunity. Whether the sex of macrophages can be causal in determining infection outcome remains unresolved.

*Drosophila melanogaster* offers an experimental system in which these limitations can be addressed directly. Its genetic tractability is unmatched when it comes to analysis of sex-specific regulation of tissue function^12–18^. Sex is determined cell-autonomously through a cell-intrinsic genetic cascade (the Sxl–tra–dsx pathway) that operates independently in each somatic cell, with no equivalent of systemic gonadal hormone signalling. As a result, sex-specific development and / or regulation of specific cells can be experimentally manipulated. Genetic tools allow for *in vivo* expression of components of the sex determination pathway, such as the female-specific isoform of *transformer* (Tra^F^), switching sex-specific regulation in one cell type while leaving the rest of the animal unchanged.

Adult *Drosophila* macrophages, known as hemocytes, are professional phagocytes that share strongly conserved features with mammalian macrophages and are central to controlling many microbial infections^19,20^. Hemocyte phagocytosis is particularly important in infections where early control of bacterial proliferation determines outcome, including by the opportunistic Gram-positive pathogen *Staphylococcus aureus*, a clinically important cause of skin, soft tissue, and bloodstream infections^21,22^. *S. aureus* frequently acquires antibiotic resistance in nosocomial settings^23^ and has a sex-biased disease burden in humans, where men are overall more prone to *S. aureus* carriage, and skin and soft tissue infections^24–29^. In vertebrates, phagocytic cells are critically important for resistance to *S. aureus* infection^22,30,31^. In flies, hemocytes act independently of the humoral immune response to control *S. aureus*^32,33^, yet how sex shapes adult hemocyte function and transcriptional responses to infection remains poorly understood^34^. One reason for this is that often, experiments that require hemocyte isolation, such as transcriptomics or functional analyses, have largely been performed in larval stages with pooled sexes^35^. A recent study has shown that hematopoiesis in the larval/pupal lymph gland is sexually dimorphic, where both systemic and cell-intrinsic mechanisms in the niche give rise to sex differences in population size, hemocyte transcription, and the structure of the developing hemocyte population^12^. Female lymph glands contain a third more hemocytes than do male lymph glands, driven by higher crystal cell numbers, and show greater hemocyte proliferation in response to bacterial challenge. These dimorphisms likely lead to sex differences in mature hemocytes, once they have dispersed from the lymph gland and adults have eclosed^12^. Because core macrophage biology is conserved across taxa, sex-dimorphic features identified in hemocytes can inform our broader understanding of how innate immune cells are shaped by sex. How sex shapes the hemocyte response to *S. aureus* infection, both at the level of transcriptional output and at the level of survival outcome, has not been resolved.

Here, we show that in a lethal *S. aureus* infection, adult male *Drosophila* succumb significantly faster than females, with faster bacterial proliferation in the initial stages of the infection. Through genetic hemocyte ablation, we determined that susceptibility to *S. aureus* is hemocyte-dependent in both sexes, with a greater contribution in females, and that females have a higher proportion of actively phagocytic hemocytes, measured by flow cytometry. To test whether the sex identity of hemocytes is itself causal, we expressed *tra^F^* specifically in hemocytes of male flies. Males with feminized hemocytes showed survival and bacterial burdens indistinguishable from females, eliminating the dimorphism. Bulk RNA sequencing of whole carcasses at 2 and 4 hours following *S. aureus* injection demonstrated a strongly sexually dimorphic response, where more genes were male-biased than female-biased, potentially a compensatory reaction to the higher bacterial burden males carry. Tissue-specific RNA sequencing on hemocytes isolated at 2 and 4 hours post-injection revealed that there are almost no detectable sex differences in induced responses to early stage *S. aureus* infection. However, male and female hemocytes have significant, sustained differences in their transcriptome, in both the absence and presence of bacterial infection. These differences include female-biased gene expression related to ROS generation and lysozyme production, which may prime female hemocytes by regulating their bactericidal capacity. Together, these results demonstrate that the cell-autonomous sexual identity of hemocytes is sufficient to drive dimorphic outcomes to *S. aureus* infection, and suggest that sex-specific phagolysosome function of innate immune cells can determine the course of infection.

## Results

### Susceptibility and resistance to *S. aureus* infection is sexually dimorphic

To test for a sex bias in *Drosophila* susceptibility to *S. aureus* infection, we injected ∼2,000 live *S. aureus* cells into four-day-old adults of both sexes carrying GFP-labelled hemocytes, *w^Dah^;HmlΔ-GAL4,UAS-GFP* (*HmlΔ>GFP*), and monitored survival. All individuals died within 72 hours (Fig. 1A), but males died significantly faster than females (Fig. 1A,B). Faster mortality can reflect reduced resistance (poorer control of bacterial growth) and/or reduced tolerance. To test resistance at the initial stages of infection, where hemocyte activity could play a significant role in limiting growth, we measured within-host bacterial load at 8 and 12 hours post-injection and quantified bacterial proliferation using negative binomial models with quadratic effect of time and a random effect intercept per fly. Load did not differ at injection (Fig. 1C). The shape of the bacterial trajectory differed between the sexes (sex × time interaction, likelihood-ratio test: χ²₂ = 6.95, *P*= 0.03; Fig. 1C), with males rising faster toward higher loads. We detected a significant difference at 8 hours post-injection, but not at 12 hours post-injection (Fig. 1C, p= 0.1), likely reflecting the increased between-individual variance in load as infection progresses, which reduces power at later timepoints^36^.

**Figure 1.**
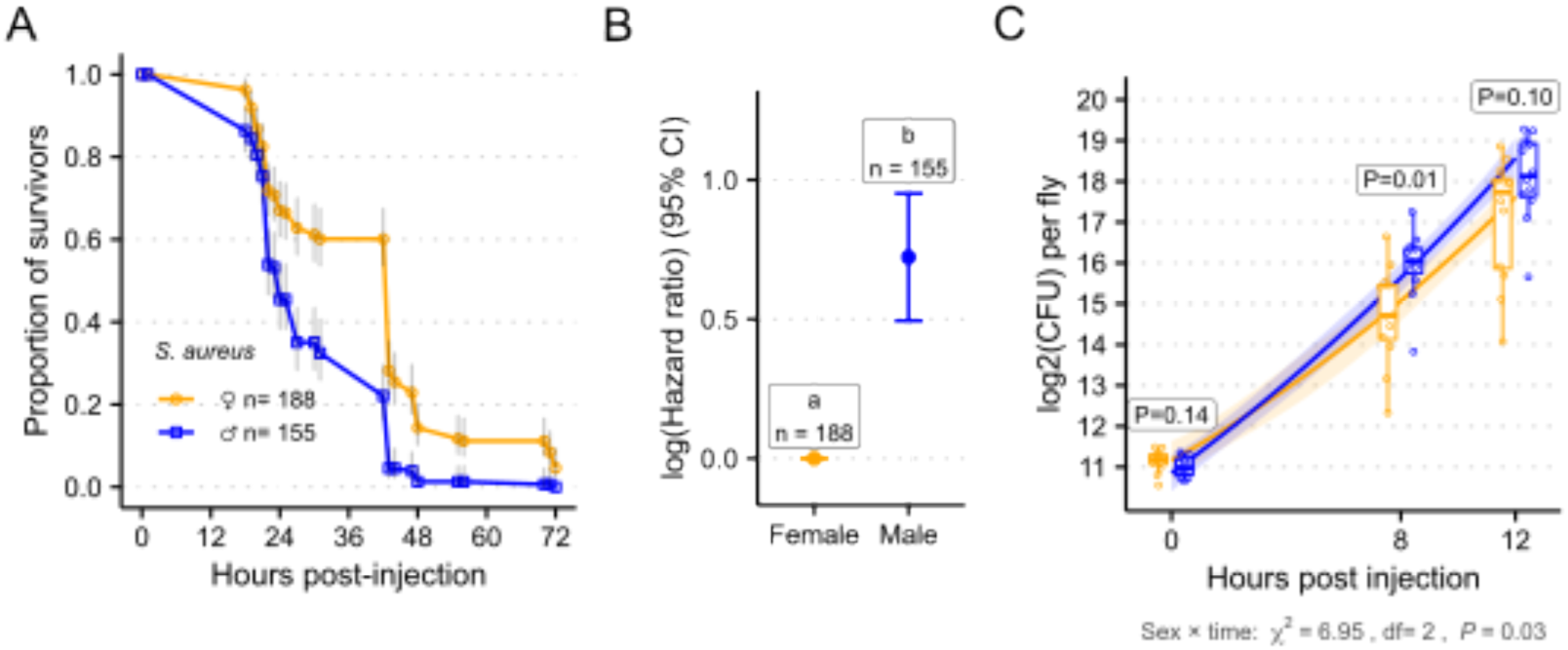
Male *Drosophila* are more susceptible to *S. aureus* infection and females better restrict bacterial proliferation. **(A)** Survival of adult *Hml>GFP* flies following injection with *S. aureus*. Points show the proportion of survivors at each scoring time, estimated by the Kaplan–Meier method; lines connect successive estimates and grey bars are 95% confidence intervals. Data pooled from two independent injection days. **(B)** Log hazard ratio (95% CI) for males relative to females, from a Cox proportional hazards model of the data in (A) with sex as a fixed effect, stratified by injection day. Females are the reference and are fixed at zero. Adding sex to a day-only model improved fit substantially (likelihood-ratio χ²₁ = 37.8, P = 7.6 x 10^10^). Shared letters indicate groups that do not differ significantly (Tukey-adjusted, α = 0.05). **(C)** Bacterial load in individual flies over the first 12 h of infection. Open circles are individual flies, boxes show the median and interquartile range with whiskers extending to 1.5× IQR, and lines with shaded ribbons are population-level fitted trajectories ± 95% CI from a negative-binomial mixed model of load against time and time², fitted separately by sex, with a random intercept per fly and an offset for plated volume and dilution. The shape of the trajectory differed between the sexes. Boxed values are per-timepoint tests for an effect of sex on load, by likelihood-ratio test against an intercept-only model of the same structure.

### Hemocytes play a significant role in resistance against *S. aureus* infection

Given the known dependency on phagocytes for host survival to *S.aureus*^32,37^, we predicted that sex differences in hemocyte function underpin the dimorphism in early bacterial proliferation and susceptibility. To test the dependency of resistance to *S. aureus* on hemocyte abundance in both sexes, we reduced the size of the hemocyte population by driving the pro-apoptotic gene, *Debcl* (*UAS-Bax*) with the hemocyte-specific, drug-inducible Geneswitch driver *HmlGS.* Adult-stage feeding of the activating drug produced the depleted genotype (‘*Hemoless’*, *HmlGS; UAS-Bax*)^38^. We focused on the initial stage of the infection, approximately the first 24 hours, before bacterial proliferation overwhelms the host. As predicted, both sexes of *Hemoless* flies showed rapid mortality upon *S. aureus* infection (Fig. 2A), confirming that hemocytes contribute importantly to resistance to *S. aureus*. Depletion increased the hazard of death substantially more in females (log-HR = 2.24, p < 0.001) than in males (log-HR = 0.75, p = 0.035). Consequently, the sex dimorphism observed in control flies (males at higher hazard; log-HR = 0.91, p = 0.02) was abolished in Hemoless flies, where the point estimate reversed (log-HR = −0.57, p = 0.12). This dependence of the sex gap on hemocyte status was confirmed by a significant sex × depletion interaction (χ²₁ = 11.8, p = 5.8 × 10⁻⁴) (Fig. 2B). This increased mortality risk in *Hemoless* flies was accompanied by an increase in bacterial proliferation of up to two orders of magnitude in both males and females, already apparent within the first 8 hours of infection (Fig. 2C).

**Figure 2.**
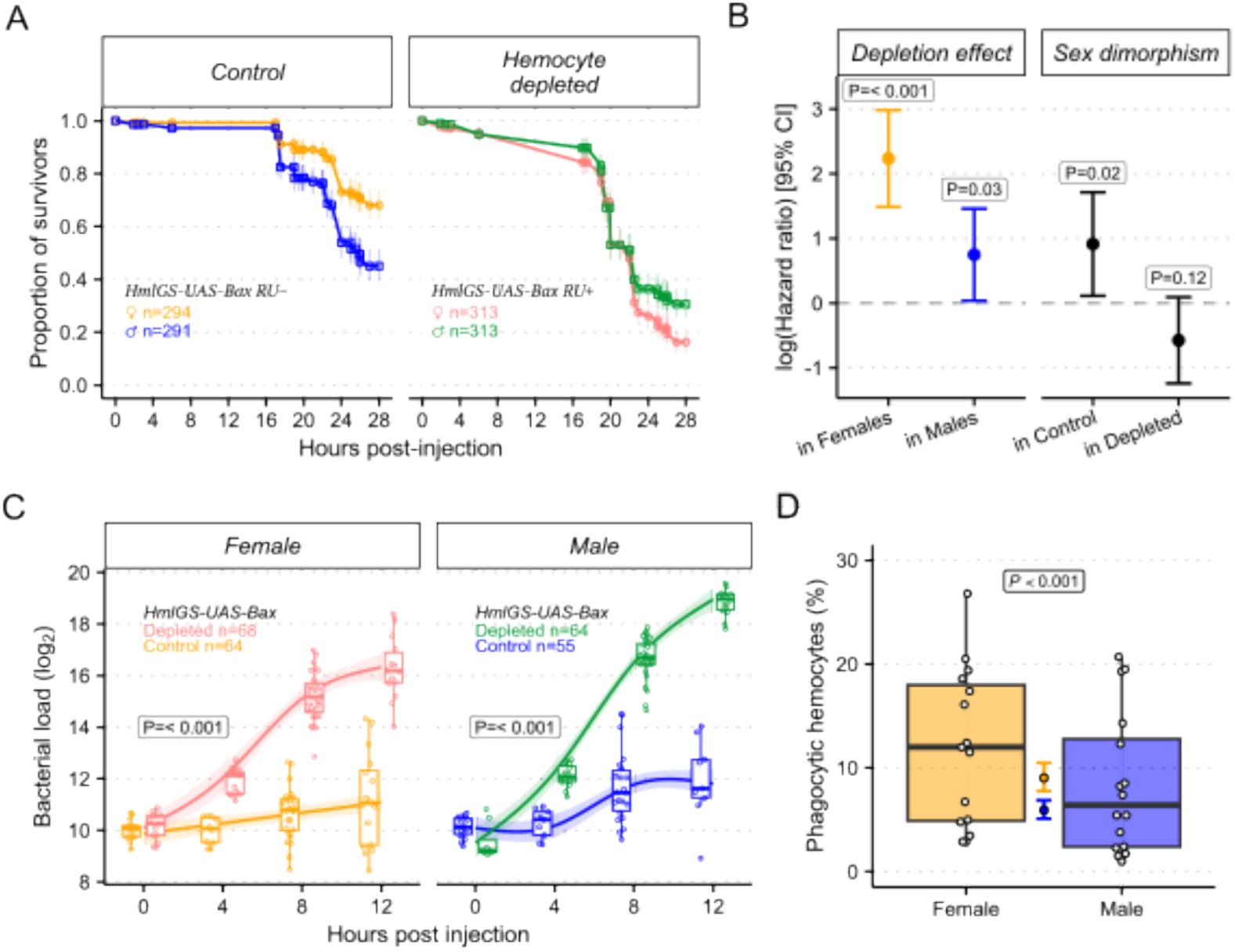
The female survival advantage against *S. aureus* is hemocyte-dependent and reflects greater per-cell phagocytic capacity. **(A)** Survival of *HmlGS>UAS-Bax* flies following injection with *S. aureus*, on control food (RU−, left) or RU486 food to induce hemocyte ablation (RU+, right). Points show the proportion of survivors at each scoring time, estimated by the Kaplan–Meier method; lines connect successive estimates and bars are 95% confidence intervals. N pooled from experiments over 5 injection days. **(B)** Log hazard ratios (95% family-wise CI) for four pre-specified contrasts, from a mixed-effects Cox model (coxme) of the data in (A) with sex × food group as a fixed effect and nested random intercepts for vial within injection day. Left, the cost of hemocyte depletion within each sex; right, the male − female difference within each food condition. Positive values indicate higher mortality. P values are single-step adjusted for the four contrasts. The effect of depletion depended on sex (sex × depletion interaction, likelihood-ratio χ²₁ = 11.8, P = 5.8 × 10⁻⁴). **(C)** Bacterial load in individual control and hemocyte-depleted flies over the first 12 h of infection, shown separately by sex. Open circles are individual flies, boxes show the median and interquartile range with whiskers extending to 1.5× IQR, and lines with shaded ribbons are fitted trajectories ± 95% CI from a generalised additive model of log₂ load against time. Boxed values are the per-sex tests of the ordered-factor difference smooth, which asks whether depletion changes the shape of the trajectory. **(D)** Percentage of GFP-positive hemocytes scoring as having engulfed fluorescent particles 4 h post-injection, measured by flow cytometry. Boxes and open circles show per-sample values; the filled point with bars is the estimated marginal mean (95% CI) from a binomial GLMM of per-cell engulfment with sex, labelling method and experimental day as fixed effects and a random intercept per sample. Data from three experimental days and two labelling methods. P value is the sex contrast on the log-odds scale.

### Females have more actively phagocytic hemocytes than do males

Having shown that hemocytes are required for the female survival advantage, we asked whether hemocytes also differ functionally between the sexes in terms of phagocytic activity. We injected flies with *S. aureus* pHRODO bioparticles, which fluoresce upon phagosomal acidification, and assayed hemocytes by flow cytometry 2 hours post-injection. Samples were run to exhaustion to recover as many hemocytes as possible and improve the estimate of absolute cell counts; while this gives only an approximate raw count, it provides a reliable basis for phagocytic index, defined as the proportion of hemocytes containing phagocytized bioparticles. Females had a larger proportion of actively phagocytosing hemocytes than males (Female − Male = 0.45 on the log-odds scale, z = 4.04, *P* < 0.001; Fig. 2D). To ask whether this translates into greater bacterial uptake at the level of the whole hemocyte population (the quantity relevant to controlling infection), we examined absolute cell counts. Neither total hemocyte number (linear model, Δ = 2728 cells, 95% CI [-1509, 6966], t = 1.32, p = 0.2, and if anything trended higher in males) nor the absolute number of engulfing cells (Δ = −111, p = 0.71) differed between the sexes. The higher female index therefore reflects a greater propensity of individual female hemocytes to engulf rather than a larger total engulfment; this absence of a sex difference in overall uptake suggests that phagocytic uptake alone is unlikely to account for the dimorphism in *S. aureus* resistance.

### Cell autonomous, sex-specific regulation of hemocytes determines outcome to S. aureus infection

Whether the genetic sexual identity of immune cells regulates their function to produce a dimorphism in resistance and survival is unknown. To test for cell autonomous sex-specific regulation in hemocytes, we expressed *transformer^Female^*(*tra^F^)*, a splicing factor which regulates female somatic sexual differentiation, specifically in hemocytes. Male cells expressing *tra^F^*adopt a feminized transcriptional profile, while all other tissues retain the endogenous sexual differentiation pathway and a male transcriptional profile. Unlike control males, males with feminized hemocytes carried the same bacterial burden as control females 8 hours post-injection (Tukey p = 0.72; Fig. 3A), whereas control males carried 2.3-fold higher loads than feminized males (Tukey p = 0.004). The survival data agreed. This experiment, performed in a different genetic background, again showed that control males were more susceptible than control females to *S. aureus* (HR = 1.68, Tukey p < 10⁻⁵), whereas males with feminized hemocytes were indistinguishable from control females (HR = 0.95, 95% CI 0.77–1.17, Tukey p = 0.88) and survived better than control males (HR = 0.57, Tukey p < 10⁻⁵; Fig. 3B, C). Together, these results suggest that cell-autonomous sex differences in hemocyte function are sufficient to account for the systemic sex differences in early bacterial load dynamics and overall survival.

**Figure 3.**
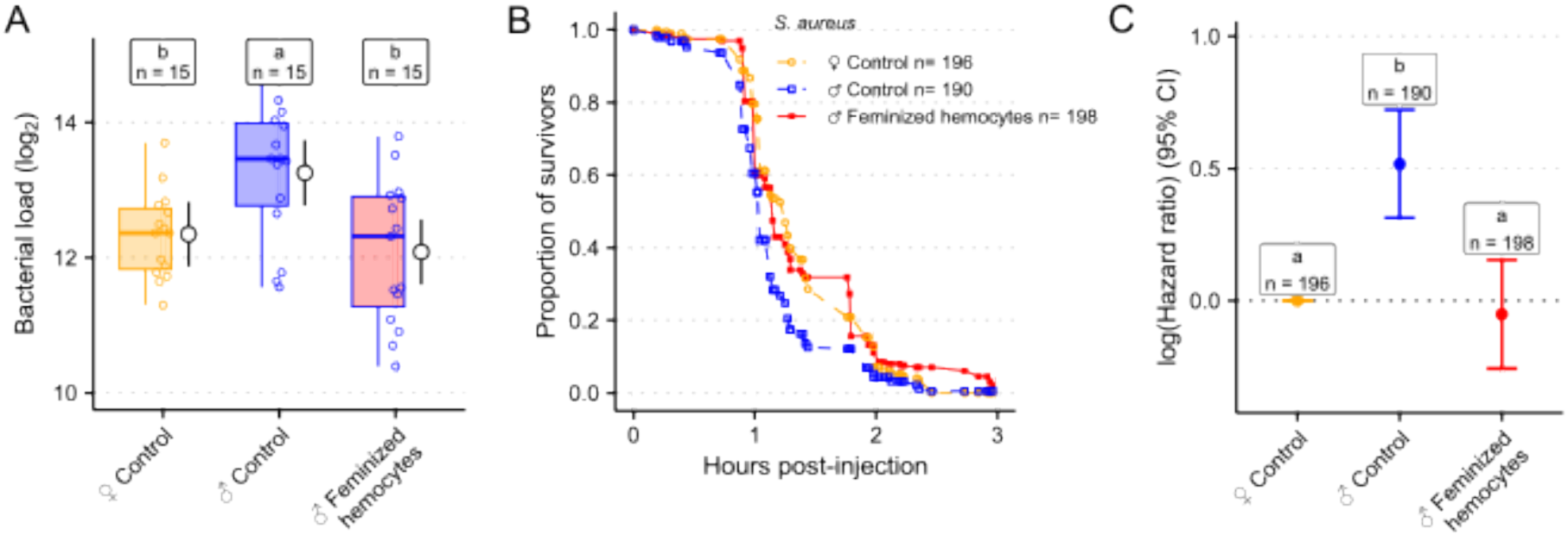
Cell-autonomous feminization of hemocytes is sufficient to confer female-like bacterial clearance and survival on male *Drosophila*. (A) Bacterial load 8 h after injection with *S. aureus* in control females, control males, and males expressing *tra^F^* specifically in hemocytes (*Hmldelta-GAL4>UAS-tra^F^*). n are pooled from two independent experiments. Boxes show the median and interquartile range, whiskers extend to 1.5× IQR, and open circles are individual flies. Black points with bars show estimated marginal means ± 95% CI from a linear model of log₂ load with genotype and experiment as fixed effects. Overall effect of genotype F₂,₄₁ = 6.58, p = 0.003. **(B)** Survival following injection with *S. aureus*. Points show the proportion of survivors at each scoring time, estimated by the Kaplan–Meier method; lines connect successive estimates. n are pooled from 4 independent experiments. **(C)** Log hazard ratios (± 95% CI) from a Cox proportional hazards model with genotype as a fixed effect, stratified by experiment. Control females are the reference and are fixed at zero. Across panels, shared letters indicate groups that do not differ significantly (Tukey-adjusted, α = 0.05).

### Systemic response to S. aureus infection differs between males and females

The result of feminization of male hemocytes suggests that sex-specific hemocyte expression underlines the sexual dimorphism of susceptibility to *S. aureus*. To understand how the immune response to *S. aureus* differs between the sexes, we performed transcriptome-wide RNA sequencing (bulk RNA-seq) on adult carcass (i.e. whole body without gonads) before infection and at 4hr post-injection, and on adult hemocytes before infection, at 2hr, and 4hr post-injection. Hemocytes were isolated from *Hmldelta>GFP* flies by FACS using an optimised extraction protocol. We first examined the systemic response by identifying genes whose change between unchallenged and infected differed between sexes (i.e. significant interaction Sex * Infection status, controlling for collection batch; Fig. 4A). The response was strongly sexually dimorphic: 429 genes showed a difference (FDR < 0.05), 248 responding more strongly in males and 181 more strongly in females. Sex-biased responders included core humoral immune genes, encoding: the AMPs *Drosomycin (Drs), Cecropin A1* and *Cecropin C*; the recognition proteins *PGRP-SC2, GNBP-like3* and *lectin-24A*; the clip-domain proteases *grass* and *SPE*; and the danger-sensing Toll protease *persephone* (*psh*), most of which are more strongly induced in males (Fig. 4B). The dimorphism extended to immune regulators, the Imd negative regulators *pirk* and *PGRP-LF* also being male-biased, indicating sex differences in the tuning, not only the magnitude, of the response.

**Figure 4.**
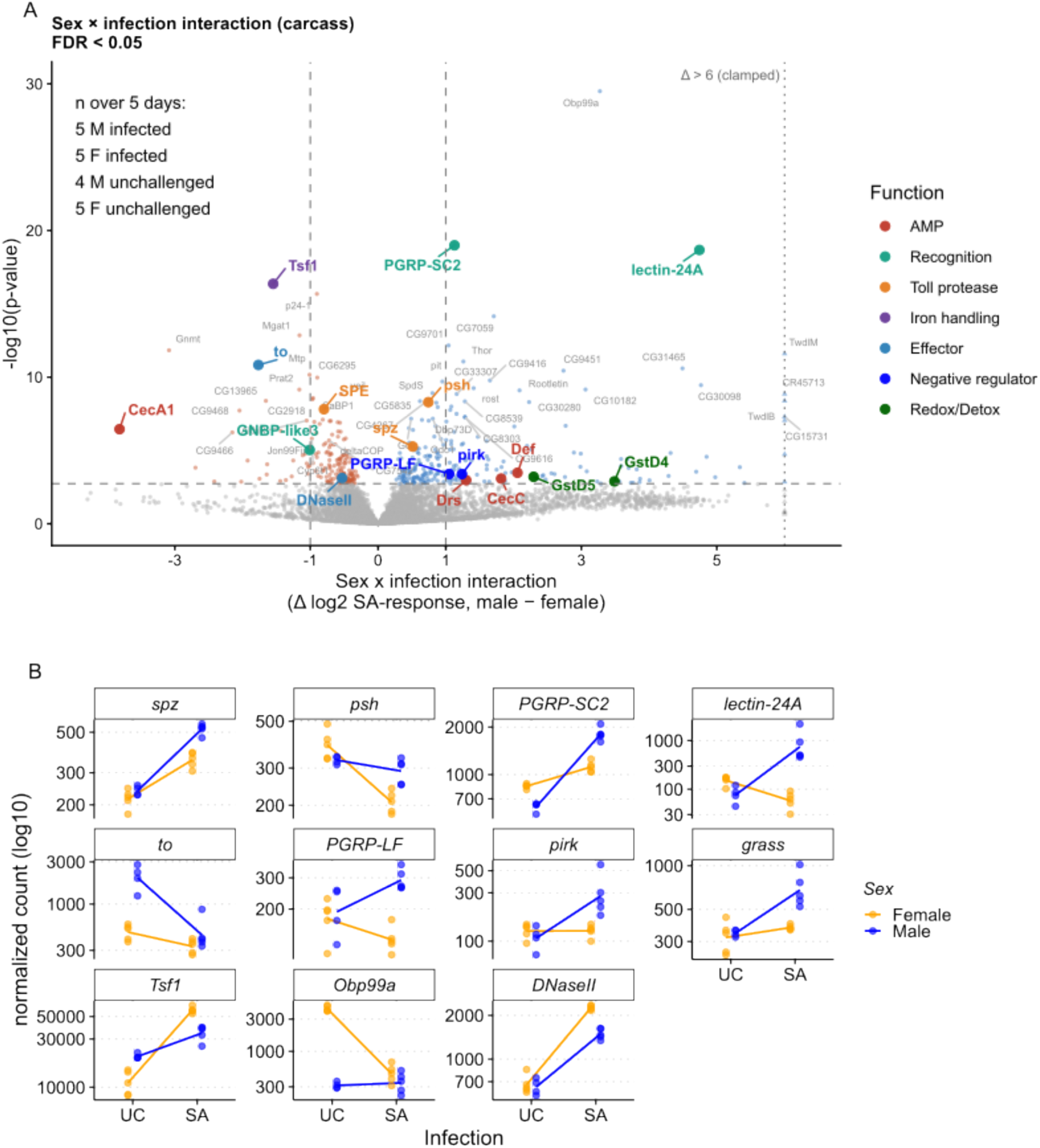
The transcriptional response to *S. aureus* infection in the carcass is sexually dimorphic. **(A)** Volcano plot of the sex × infection interaction in carcass tissue. Each point is a gene; the x-axis is the interaction coefficient, so positive values indicate a stronger infection response in males. Coloured points are genes of interest grouped by function; grey labels are the 30 genes per direction with the lowest adjusted p-values, excluding the highlighted set. Dashed horizontal line, FDR = 5%; dashed vertical lines, |log₂ fold-change| = 1. Interaction coefficients beyond ±6 are clamped to the axis limit and drawn as triangles. **(B)** Normalised counts for selected sex-dimorphic genes in unchallenged (UC) and *S. aureus*-infected (SA) flies. Points are individual samples and lines connect group means; y-axis log₁₀ with a free range per gene.

Several of these genes have independent support: *Drs, Tsf1, Defensin* (*Def*) and *takeout* (*to*) were reported as showing sexually-dimorphic expression following *P. rettgeri* infection, where the dimorphism was localised specifically to the persephone branch of Toll^36^, the same branch our *psh* result implicates.

The predominance of male-biased induction is notable given that males are the more susceptible sex to *S. aureus*^36^. A stronger systemic response in males therefore does not confer protection: it may indicate that this humoral response is ineffective against *S. aureus*, or that it is a compensatory reaction to the higher bacterial burden males carry. Consistent with the response being insufficient regardless of magnitude, *S. aureus* is lethal a few hours post-injection. In contrast, two effectors were more strongly induced in females, the iron-sequestration gene *Transferrin 1* (*Tsf1*) and the phagolysosomal nuclease *DNase II*, which degrades ingested bacterial DNA during phagocytic clearance, making them candidate contributors to the female advantage.

### Hemocytes selectively upregulate ribosome-biogenesis genes alongside the shared humoral response

Having established that the systemic response is sexually dimorphic, we investigated which components of the transcriptional infection response are tissue-specific. We compared the *S. aureus*-induced log₂ fold-changes (SA vs. UC) between isolated hemocytes and the carcass, controlling for sex as a covariate (Fig. 5A).

**Figure 5.**
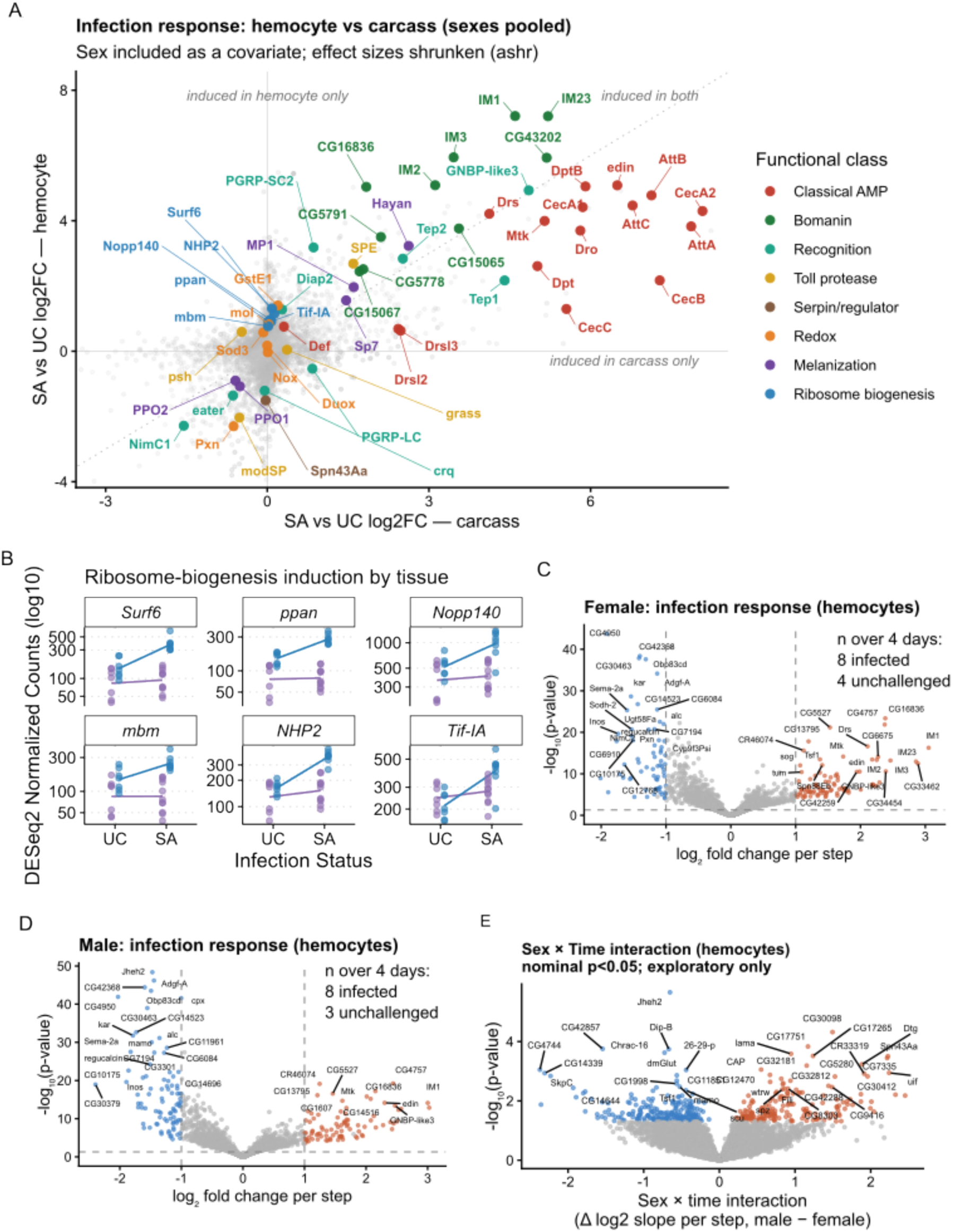
Hemocytes mount a distinct transcriptional response to *S. aureus*, and this response does not differ detectably between the sexes. **(A)** Infection response in hemocytes plotted against the response in carcass, sexes pooled. Each point is a gene; axes are ashr-shrunken log₂ fold-changes for infected (4 h) versus unchallenged flies, estimated within each tissue. Grey point opacity scales with the significance of the tissue × infection interaction. Coloured points are genes of interest grouped by functional class; the dotted diagonal marks equal induction in both tissues. **(B)** Normalised counts for ribosome-biogenesis genes in unchallenged (UC) and infected (SA) carcass and hemocyte samples. Points are individual samples and lines connect group means; y-axis log₁₀ with a free range per gene. **(C, D)** Volcano plots of the hemocyte infection response over time in females (C) and males (D). The x-axis is the log₂ fold-change per timepoint step; coloured points pass FDR < 0.05 with |log₂ fold-change| > 1. Dashed lines mark p = 0.05 and |log₂ fold-change| = 1. **(E)** Volcano plot of the sex × time interaction in hemocytes, that is, the difference between male and female response slopes. Coefficients are unshrunken, so that effect sizes and Wald p-values remain on the same footing; coloured points are nominally significant (p < 0.05). Only one gene passed FDR < 0.05, so this panel is exploratory and no gene is claimed as sex-dimorphic.

At the level of direct immune effectors, the two tissue compartments responded concordantly. Classical antimicrobial peptides (*Cecropins*, *Attacins*, *Diptericins*, *Drosomycin*, *Metchnikowin*) and Bomanins were strongly induced in both settings, clustering near the diagonal line of equal induction (Fig. 5A). While shrunken fold-change estimates suggested subtle tissue biases, with Bomanins leaning slightly toward hemocytes and classical AMPs toward the carcass, underlying normalised counts revealed that this largely reflected baseline differences rather than divergent induction dynamics (Fig. S1 & S2). Both effector classes rose strongly with infection in each compartment, though the carcass, containing the fat body, remains the dominant absolute pool of systemic transcripts owing to its mass. Thus, the inducible peptide-effector repertoire is broadly shared between the humoral and cellular compartments rather than partitioned between them.

By contrast, hemocytes mounted a prominent cellular activation response that contrasted with the stable baseline of the carcass. This signature was dominated by genes essential for ribosome biogenesis, nucleolar scaffolding and rRNA processing, including *Surf6*, *ppan*, *Nopp140*, *NHP2* and the *Pol I* transcription factor *Tif-IA* (Fig. 5A, B). While these biogenesis factors maintained comparable expression baselines between tissues in uninfected flies, they were robustly upregulated in hemocytes upon infection while showing little or no induction in the bulk carcass (Fig. 5B). This is not a hemocyte-private effector programme: the preferential upregulation of translational machinery instead indicates a cellular activation response specific to the hemocyte compartment.

Finally, the reactive oxygen species (ROS)-generating oxidases Duox and Nox were not appreciably induced by infection in either tissue compartment. This lack of transcriptional upregulation suggests that oxidative defence may rely on baseline expression or post-translational activation rather than acute *de novo* transcription, leaving the upregulation of ribosome-biogenesis factors as the most prominent active metabolic investment at this time point.

### The temporal kinetics of the hemocyte response do not differ detectably between the sexes

To characterise the temporal dynamics of the cellular immune response, we modelled the hemocyte transcriptional trajectory across a post-infection time course (uninfected control, 2 hours, and 4 hours), parameterising infection duration as a continuous step-variable. Analysed separately, both male and female hemocytes mounted a clear kinetic response of comparable scale. In females, 158 genes were responsive per temporal step (91 upregulated, 67 downregulated; FDR < 0.05, |log₂FC| > 1; Fig. 5C), and a similar number was observed in males, with 175 genes altered per time step (87 upregulated, 88 downregulated; Fig. 5D). The two responsive gene sets overlapped substantially (109 genes shared, 69% of the female set and 62% of the male set), consistent with a conserved core programme of cellular activation, though we note that this list overlap understates the true similarity, since genes with comparable trajectories can fall on opposite sides of a significance threshold.

We next tested explicitly for sexual dimorphism in these trajectories using a genome-wide sex × time interaction model. We found no evidence for a sex-dependent hemocyte response: only a single gene, *Jheh2* (encoding a juvenile hormone epoxide hydrolase), passed the false discovery rate threshold (FDR < 0.05; Fig. 5E). Furthermore, its underlying counts revealed a shared downregulation in both sexes, differing only in slope from a higher male baseline, rather than a sex-private response.

To determine whether this null result reflected a genuine absence of dimorphism rather than limited detection power, we estimated the proportion of true null hypotheses (π₀) across the whole transcriptome and compared it with the equivalent quantity from the carcass. In hemocytes, π₀ = 0.97 for the sex × time interaction, whereas in carcass π₀ = 0.77 for the sex × infection interaction, computed by the same method on a dataset of comparable size and replication. At least 23% of carcass genes therefore carry a sex-dependent infection response, against at most a few per cent in hemocytes. This suggests a roughly sevenfold difference in the extent of dimorphic signal between the two compartments.

Thus, the temporal kinetics of the hemocyte infection response do not differ detectably between the sexes, though we note that this analysis rests on three to four biological replicates per sex per timepoint and cannot exclude effects smaller than that sample size can resolve. Because our model statistically partitions the constitutive sex difference (sex main effect) from the sex-specific response (sex × time), this result localises any hemocyte dimorphism to sustained, baseline differences rather than to the dynamics of the acute induced response, which we examine next.

### Hemocytes show a constitutive, sexually dimorphic redox, lysozyme and protease-inhibitor profile

Having found no sex difference in the dynamics of the hemocyte infection response, we asked whether the sexes differ in their constitutive hemocyte transcriptome. Modelling the sex difference as a sustained offset across the time course (Methods), we found that constitutive differences were extensive: 441 genes were female-biased and 1,594 male-biased (FDR < 0.05, |log₂ fold-change| > 1; Fig. 6A). Much of this difference reflected organismal rather than immune sexual identity: among the most strongly male-biased genes are the dosage-compensation lncRNA *roX1*, the male-specific serine protease homologues *scpr-A*–*C*, and the sperm-leucylaminopeptidase *S-Lap5*, none of which has a known hemocyte function. We therefore focused on functional categories with a plausible role in bacterial killing, rather than on the global gene list. Three such categories showed a clear sex bias.

**Figure 6.**
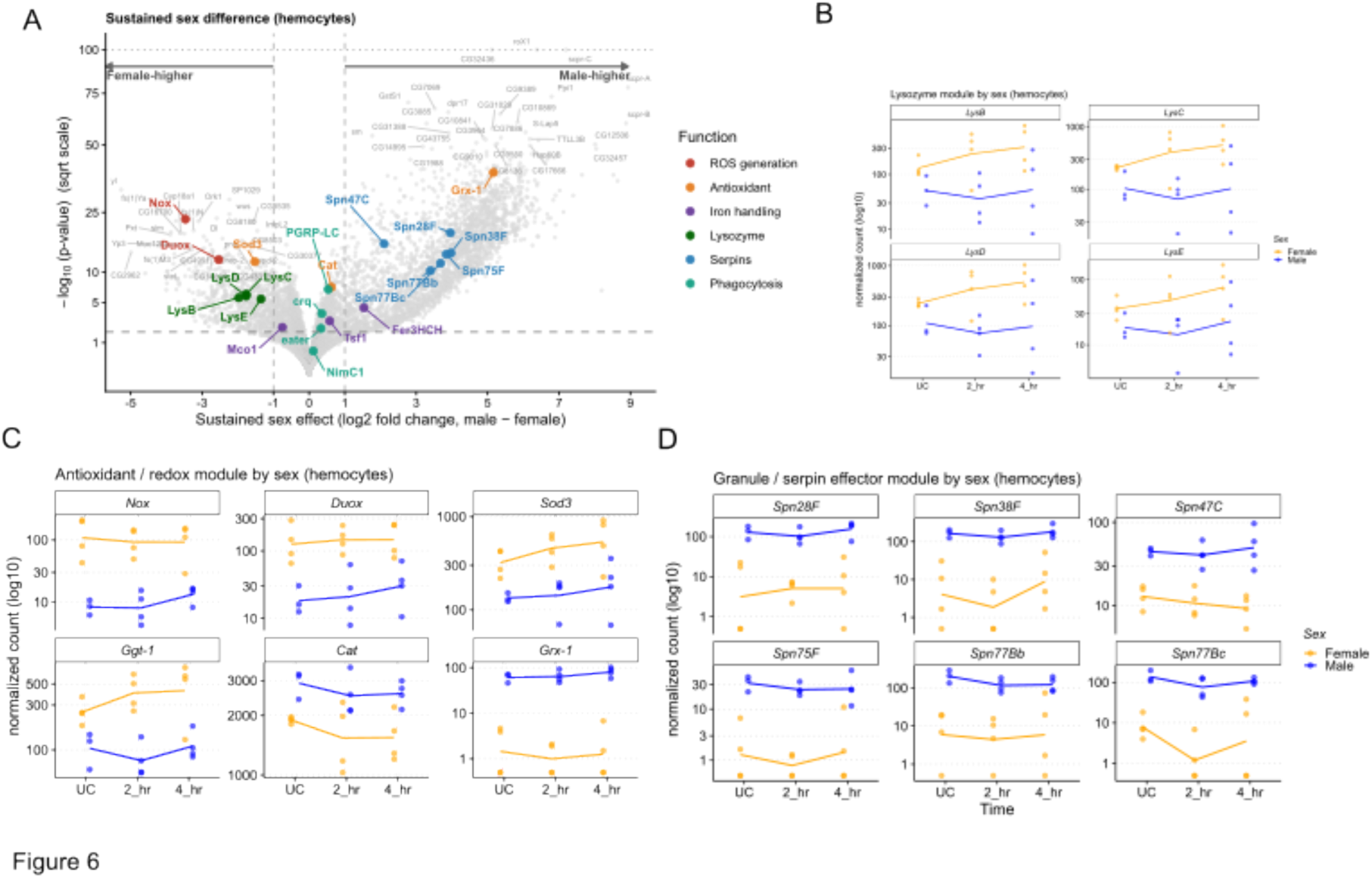
Hemocytes carry a sustained, infection-independent sex difference in redox and serpin gene expression. **(A)** Volcano plot of the sustained sex difference in hemocytes, estimated as a constant offset between sexes across the infection time course. Each point is a gene; positive values indicate higher expression in males. Coloured points are genes of interest grouped by function; grey labels are the 30 genes per direction with the lowest adjusted p-values, excluding the highlighted set. Dashed horizontal line, FDR = 5%; dashed vertical lines, |log₂ fold-change| = 1. The y-axis is on a square-root scale, and p-values below 10⁻¹⁰⁰ are clamped to that value (dotted line). **(B)** Normalised counts for the lysozyme module in unchallenged (UC) and infected hemocytes at 2 h and 4 h. Points are individual samples and lines connect group means; y-axis log₁₀ with a free range per gene. **(C)** Normalised counts for the antioxidant/redox module, plotted as in (B). **(D)** Normalised counts for the granule/serpin effector module, plotted as in (B).

First, and most relevant to bacterial killing, the ROS-generating NADPH oxidase *Nox* was expressed 11-fold higher in female hemocytes (padj = 2.6 × 10⁻²²), with *Duox* showing the same bias at 5.8-fold (padj = 2.4 × 10⁻¹²; Fig. 6A, C). This bias was specific to ROS generation rather than reflecting a general redox shift: the surrounding antioxidant machinery did not move in concert, with *Sod3* female-higher but *Cat* and the thiol buffer *Grx-1* male-higher (Fig. 6C). Females therefore do not simply have greater global antioxidant capacity, but specifically elevated ROS-generating capacity.

Second, females also constitutively expressed higher levels of the lysozyme cluster (*LysB*–*E*, 2.5– to 3.9-fold, padj < 2 × 10⁻⁵ for all four; Fig. 6A, B). These are abundant transcripts in female hemocytes specifically: *LysC* and *LysD* rank above the 90th percentile of expressed genes in females but around the 80th in males, and *LysB* falls from the top decile in females to mid-range in males. Together with the ROS result, this indicates that two components of phagolysosomal bacterial killing are constitutively elevated in female hemocytes.

Conversely, a coherent set of six serpins (*Spn28F*, *Spn38F*, *Spn47C*, *Spn75F*, *Spn77Bb*, *Spn77Bc*) was strongly male-biased, with sustained differences of 2.1 to 4.0 log₂ units (padj < 1.5 × 10⁻¹⁰ for all six; Fig. 6A, D), indicating a sex difference in the constitutive proteolytic-inhibitor balance of hemocytes. Phagocytic receptors (*eater*, *NimC1*, *crq*, *PGRP-LC*) showed no consistent female bias, consistent with our functional finding that total bacterial uptake does not differ strongly between the sexes (Fig. 2D).

These constitutive differences identify elevated female ROS-generating capacity, alongside elevated lysozyme expression, as candidate mechanisms for the female survival advantage.

## Discussion

Differences in immunity between males and females are striking, but historically neglected^34,39^. Although sex-stratified immunology research is gaining traction^40,41^, substantial knowledge gaps were called into sharp focus during the recent global pandemic where men were at significantly higher mortality risk than were women^42^. Research into immune dimorphism is also critical to understanding infection dynamics and life history in animal populations, including insects^34,43^. Although immune cells are likely to play a significant role in dimorphic responses to infection, there are scant causal data on sex differences in innate immune cell responses to *in vivo* infection^9^, including dynamic behaviours like phagocytosis, and determinants of microbicidal function, in any model system. Here, in a *Drosophila* – bacterial infection model, we show that survival to infection by gram-negative *S. aureus* is female-biased. Analysis of bacterial load in early stages of infection suggested that females are better able to limit proliferation of *S. aureus*, and we demonstrated via genetic ablation that hemocytes are essential for this early control. This places hemocytes in a similarly central position to phagocytes in *S. aureus* bloodstream and wound infections in mammals, where neutrophils are the dominant early protective cells, and macrophages are important for containment and clearance over time^37,44^. Across *S. aureus* infection models in mice and zebrafish, innate phagocytes are causal determinants of early outcome: depleting or impairing neutrophil or monocyte effector functions increases bacterial burden and worsens survival^30,31,45–48^, similar to our observations in hemocyte-ablated flies.

In *Drosophila*, the sexes may differently regulate hemocyte function non-cell autonomously through systemic immune responses to *S. aureus* infection or via hormonal control. Alternatively, the cell-autonomous sex identity of hemocytes may determine their activity. To distinguish between these two possibilities, we took advantage of the ability to switch sex-specific transcriptional regulation of hemocytes, by expressing the female form of the sex-determining splicing factor, *tra^F^*, in mature hemocytes. This not only reduced the early bacterial burden in males to levels comparable to that of females, but it also lowered male mortality risk to match females. This points to cell-autonomous, *tra*-mediated regulation of hemocyte resistance mechanisms during *S. aureus* infection, where hemocyte dynamics at early timepoints can determine overall mortality risk.

Our data synergise well with that presented in a recent study of sex-specific control of hematopoiesis in the larval/pupal lymph gland^12^. Using a similar *tra*-misexpression approach under the control of lymph gland-specific drivers, this study showed that cell-autonomous, sex-specific regulation in the niche gives rise to sex differences in hematopoiesis, transcription, and the overall structure of the developing hemocyte population. This study also identified insulin signalling regulation of the niche, suggesting that organismal sex can also influence hemocyte development. An interesting outcome from this study was that females produce more hemocytes in the lymph gland overall, driven primarily by higher numbers of crystal cells, where sex differences in plasmatocyte number were not detected. Consistently, we did not detect sex differences in overall adult plasmatocyte number. To note, the driver we used to express GFP for flow cytometry and to mis-express *tra^F^* in hemocytes (*HmlΔ-Gal4*) is only expressed in mature plasmatocytes^49^, and thus, is not expected to influence hematopoiesis. While lineage tracing suggests approximately 60% of adult hemocytes derive from embryonic hematopoiesis^50^, both waves of blood cell development contribute to determining overall adult hemocyte number, population structure, and function. Sex differences in embryonic hematopoiesis are so far unexplored, thus the relative contribution of both waves of hematopoiesis to dimorphisms in adult populations is still an open question. Although we did not detect differences in hemocyte number, we did find a difference in phagocytic index, where female hemocytes were more likely to phagocytose injected *S. aureus* particles. Although this difference was subtle, it is likely to be underestimated, and could make some contribution to more efficient control of bacterial proliferation in females.

Our data suggest that cell-intrinsic hemocyte dimorphisms include the baseline sex-biased transcriptional regulation of bactericidal functions like ROS generation, lysozyme production and Serpin expression. We did not detect sex differences in the acute hemocyte response to *S. aureus* infection: in fact, hemocytes from males and females showed very similar transcriptional early response profiles, with substantial overlap between core response gene sets, and without any detectable dimorphic responses. The sex differences that we detected were sustained – that is, they were different both at baseline, and during infection. This suggests that female hemocytes, which express higher levels of ROS-producing enzymes and antimicrobial lysozymes, are poised to control bacterial proliferation more efficiently compared to male cells. In fast-acting mechanisms such as phagocytosis and bacterial destruction in the phagolysosome, baseline transcription will determine efficiency. These data are intriguing given that production of lysozymes has not yet been explicitly linked to hemocyte function in flies, despite their known role in mammals, where innate cell effector mechanisms such as NET formation, and ROS production and AMP generation (including lysozymes) support *S. aureus* clearance^22^. Thus, sexually dimorphic phagocyte functions, such as the mechanisms we have uncovered here, have the potential to influence the sex differences in *S. aureus* infection outcome observed in mammalian and clinical data. In humans, men are more prone to skin and soft tissue infections by *S. aureus*, an infection stage that is controlled by phagocytes. Although dimorphic innate cell responses specifically to *S. aureus* infection have not been investigated in mammals to our knowledge, recent studies on sex differences in rodent and human peripheral blood have demonstrated that macrophages stand out among leukocytes as having sexually dimorphic transcriptomes^5–7^. IFN signalling shows sustained sex-bias at baseline and following stimulation, where female macrophages appear to be in a ‘poised’ state for IFN-mediated responses to viral infection^7^. This suggests that innate cell antimicrobial readiness may be a broadly conserved feature of immune dimorphism.

Baseline sex differences in transcriptional regulation in macrophages, such as the signatures that we detected, could reflect other functional constraints that differ between males and females, in addition to infection responses. By definition, the sexes have different reproductive biology, which imposes distinct requirements in terms of physiology, metabolism, and behaviour, and distinct selective pressures on males and females. The impact of sex differences are highly tissue– and context-specific, but likely have their root in sex-specific fitness optima, where most, if not all, tissues show sex-biased cellular regulation and metabolism. For example, *Drosophila* gut cells respond to nutrients with sex-biased growth and metabolism to maximise egg or sperm production^13,14^. Consequences of metabolic dimorphism include differences in responses to intestinal infection^51^, and the rate of intestinal ageing^16,52^. Sex differences in immune cells may also be implicit in metabolic fine-tuning to enhance reproductive fitness, or directly selected for, if one sex is particularly vulnerable or more frequently exposed to pathogenic challenge because of their distinct ecologies^53^. Macrophages are multitasking cells, involved in homeostatic functions such as efferocytosis and tissue remodelling, as well as host defence^54^; therefore, the drivers and impacts of sex-biased macrophage phenotypes are likely diverse and difficult to disentangle. Sex differences in macrophage behaviour can even shape developmental processes. For example, in the mammalian brain, females have more phagocytic microglia than males, contributing to differences in axonal pruning that influence brain development and subsequent juvenile behaviour^55^. Given the diverse roles performed by macrophages, infection control may only be one aspect of their dimorphic function.

Sex-biased macrophage function has the potential to shape host defence, development, ageing, and disease progression, and thus, there is a clear incentive for understanding underlying sex-specific regulation of innate cells. While some sex-specific macrophage functions are likely to be particular to species ecology, other dimorphisms may be deeply conserved, such as responses to bacterial challenge, as described here for *S. aureus* infection which is female-biased from flies to humans. *Drosophila* offers the genetic tools to unpick the basic biology of sexually dimorphic macrophage development and function, and its impact on systemic physiology. Our study has uncovered that cell-autonomous, sex-specific regulation of macrophages directly determines bacterial load and survival to acute bacterial infection, and that sex-biased transcriptional regulation favours a pathogen-ready state in female cells.

## Methods

### *D. melanogaster* stocks and husbandry

Flies were maintained at 25 °C in 60–70% humidity on a 12:12 light:dark cycle. The following *Drosophila* stocks were used: *w^Dah^*;*HmlΔ-GAL4,UAS-GFP* (Bloomington Drosophila Stock Center, BDSC #30140, backcrossed for 7 generations into the outbred line, *w^Dahomey^*)^49^; *Hml-GS*; *UAS-Debcl (UAS-Bax;* BDSC #58357); *UAS-traF* (BDSC #4590). Transgenic lines were backcrossed into the outbred line, *wDah*, for 7 generations before use. All lines were expanded in a standardised, density-controlled manner in bottles containing standard sucrose and yeast (SYA) food (Bloomington Drosophila Stock Center, 2021), repeated for a minimum of four generations to control for transgenerational effects of crowding. All experiments were conducted across at least three replicates, which implies different days, different cohorts, and different bacterial preparations.

### Systemic infection with *S. aureus*

The Gram-positive bacteria, *Staphylococcus aureus* (PIG1 strain)^56^, was streaked onto LB agar plates from frozen 25% glycerol stocks and incubated at 37 °C. A single colony was suspended in Luria-Bertani broth and incubated overnight at 37 °C. The resulting culture was pelleted and resuspended in PBS to an optical density at 600 nm (OD_600_) of 0.25 Four day post-eclosion flies were anaesthetised with CO_2_ and injected in the abdomen with 23 nL of bacterial suspension using a Nanoject III microinjector (Drummond)^57^, corresponding to a dose of approximately 2000 viable *S. aureus* bacteria per fly. For the control treatment, flies were injected with 23 nL sterile PBS into their abdomen. Flies were observed after injection to ensure recovery, and stored in single sex vials containing SYA food in groups of 15 individuals. Survival was monitored at least 10 times over the course of 24-72 hours post-injection. Survival responses were analysed using the R package, “survival”^58,59^. Host survival differences were analysed using a cox proportional hazards model with hemocyte depletion or sex as the main effects and date and vial replicate as random effects; hazard ratios were extracted from this model.

### Bioparticle injection for phagocytosis analysis

23 nL (1 mg/mL) of pHrodo red *S. aureus* bioparticles (ThermoFisher A10010) was injected into the abdomen of 4 day old, adult *Drosophila* anaesthetised with CO_2_. Recovery was observed after injection, and flies were incubated for 2 hours at 25°C. Hemocytes were extracted for flow cytometry analysis as described below.

### Bacterial load measurement – colony forming unit (CFU) estimation [27]

Individual flies were homogenised in 500 mL PBS (Merck 66062) with two sterile glass beads in a TissueLyser II (Qiagen) at 25 Hz for 45 seconds. Homogenate was diluted 1:5, 1:25, and 1:125 with PBS and 5 μL of each dilution was plated in duplicate, including undiluted, on LB agar plates. Plates were stored at 25°C overnight, then transferred to 37°C for 2-5 hours, until colonies were visible. Plate images were captured, and bacterial colonies were counted manually, blind. Data were tested with a linear mixed-effect model in R using the lmer function from the package “lme4”^60^, with technical replicate as a mixed effect.

### Adult hemocyte isolation

Pools of 30 four-day-old individuals were anaesthetised on ice before being crushed with a pestle in 500 μL ice-cold PBS. Homogenate was then transferred to a 40 μm filter atop a 50 mL Falcon tube. An additional 500 μL of PBS was used to rinse the tube and filter. The flow through was transferred to a clean 1.5 mL tube and centrifuged at 4°C for 3 minutes at 1200 g. The samples were then washed three times by removing the supernatant, adding 500 μL ice-cold Schneider’s medium (Sigma-Aldrich), and centrifuging at 4°C for 3 minutes at 1200 g.

### Larval hemocyte isolation

Male and female L3 *HmlΔ-GAL4, UAS-GFP* larvae were sexed by the size of the gonadal discs^61^. Larvae were pooled by sex into groups of 10 and placed on a chilled cavity slide on ice in 100 μL cold PBS (Merck 66062). The cuticle of each larva was pierced, taking care to avoid the gut. Larvae were bled for 5 minutes, and PBS / hemolymph was transferred to a 1.5 mL tube in ice. The slide was rinsed with 100 μL PBS, which was added to the sample.

### Flow cytometry

Adult *HmlΔ-GAL4, UAS-GFP* hemocyte samples (prepared as described above) were transferred to a 50 μm filter and collected in a clean 5 mL round bottom tube, immediately prior to running on the flow cytometer (BD LSRFortessa). Flow cytometry data were analysed with FlowJo v10.8.1. Cells were gated to exclude debris and doublets by first gating based on FSC-A versus SSC-A and excluding cells with low values of both. Single cells were next gated by viewing FSC-W versus FSC-A and selecting cells under 125K and above 50K. Hemocytes were identified by fluorescence intensity on the FITC channel, and bioparticles by fluorescence intensity on the PE-142 Texas Red channel. Labelled hemocytes that contained bioparticles were identified as phagocytic hemocytes. Phagocytic index was tested using a binomial generalised linear mixed-effects model with sex as a fixed effect, using the R package “lme4”^60^. Data were visualised using the R package “ggplot2”^62^.

### RNA-extraction and library preparation

Adult and larval *HmlΔ-GAL4, UAS-GFP* hemocytes were isolated as described above, and sorted by FACS directly into 750 μL TRIzol. RNA was extracted from sorted hemocytes using a Direct-zol kit (Zymo) following manufacturer’s guidelines. Quality of RNA was measured using a 2100 Bioanalyzer (Agilent Technologies) and RNA quantities were measured with a Qubit III using an RNA HS assay (Thermo Fisher). RNA-sequencing libraries were then prepared using the NEBNext Ultra II RNA Library Prep Kit for Illumina with the NEBNext Poly(A) mRNA Magnetic Isolation Module (New England BioLabs).

### RNA-sequencing

All sequencing and mapping were performed by Edinburgh Genomics. Libraries were sequenced on a NovaSeq SP 150PE cycle setup to yield approximately 375M read pairs. Reads were trimmed using Cutadapt (version cutadapt-1.18-venv). Reads were trimmed for quality at the 3’ end using a quality threshold of 30 and for adapter sequences of the TruSeq stranded mRNA kit (AGATCGGAAGAGC). Reads after trimming were required to have a minimum length of 50. Reads were aligned to the reference genome, *Drosophila melanogaster* BDGP6, using STAR2 (version 2.7.3a) specifying paired-end reads and the option – outSAMtype BAM Unsorted. All other parameters were left at default. Reads were assigned to features of type ‘exon’ in the input annotation grouped by gene_id in the reference genome using featureCounts3 (version 1.5.1). Genes with biotype rRNA or Mt_tRNA were removed prior to counting. featureCounts assigns counts on a ‘fragment’ basis as opposed to individual reads such that a fragment is counted where one or both of its reads are aligned and associated with the specified features. Strandedness was set to ‘reverse’ and a minimum alignment quality of 10 was specified. In addition to the counts matrix used in downstream differential analysis, a matrix of Fragments Per Kilobase of transcript per Million mapped reads (FPKM) values was generated, using the rpkm() function of edgeR4 (version 3.28.1) and normalised effective library sizes. Gene lengths for the FPKM calculation were the number of bases in the exons of each gene (only counting bases once where they occur in multiple exon annotations). FPKM values may be useful in analyses requiring accurate gene-gene (as opposed to sample-sample) comparisons. Gene names and other fields were derived from input annotation and added to the count/expression matrices.

### RNA-sequencing analysis

Differential expression was assessed on raw fragment counts with DESeq2 size factors, which correct for sequencing depth and library composition. Length-normalised units (FPKM, TPM) were not used for these analyses, since gene length is constant across samples and does not affect within-gene comparisons; FPKM values are provided for cross-gene comparisons.

One collection day (13/7) separated clearly from the remaining batches on principal component analysis of the hemocyte samples and was excluded from the hemocyte time-course and tissue-comparison analyses. The same batch was not an outlier in the carcass samples and was retained there, giving five collection days for the carcass analysis and four for the hemocyte analyses.

Throughout, ashr-shrunken fold-changes are used for effect-size display and ranking, since shrinkage moves imprecisely estimated effects toward zero in proportion to their standard error and prevents low-count genes with large apparent fold-changes from dominating. Significance testing was performed on unshrunken maximum-likelihood estimates and their Wald p-values, so that reported effect sizes and p-values remain on the same footing.

#### Sex-dependence of the carcass infection response

Carcass samples from adult flies were sequenced from unchallenged flies and at 4 h post-injection with *S. aureus*, in both sexes, leaving 11,372 genes. Counts were modelled in DESeq2 (v. 1. 48) with the design ∼ Date + Sex + Infected + Sex:Infected, with collection date as a fixed blocking factor. Sexual dimorphism in the infection response was tested through the Sex:Infected interaction term, which contrasts the male and female log₂ responses to infection. Genes were called dimorphic at FDR < 0.05, with directionality taken from the sign of the maximum-likelihood coefficient. Interaction coefficients exceeding ±6 log₂ units were clamped to the axis limit for display.

#### Sex-dependence of the hemocyte infection response

Hemocyte samples from adult flies were sequenced at three timepoints (unchallenged, 2 h and 4 h post-injection) in both sexes, leaving 10,745 genes. Counts were modelled with the design ∼ Date + Sex + Time + Sex:Time, with collection date as a fixed blocking factor. Time was coded linearly as a numeric covariate (unchallenged = 0, 2 h = 1, 4 h = 2), so that each coefficient represents the change in expression per step. This assumes a constant per-step change on the log scale, that is, that the shift from unchallenged to 2 h equals that from 2 h to 4 h; unchallenged samples serve as the time-zero baseline. The female response slope was taken from the Time coefficient and the male slope from the sum of the Time and Sex:Time coefficients. Genes were called responsive at FDR < 0.05 with |log₂ fold-change| > 1. Sexual dimorphism in the response was tested through the Sex:Time interaction term.

#### Genome-wide extent of sex-dependent signal

To assess how much sex-dependent signal was present beyond the genes passing FDR correction, the proportion of true null hypotheses (π₀) was estimated from the distribution of interaction p-values with the qvalue package (v. 2.4). The estimator is conservatively biased upward, so 1 − π₀ provides a lower bound on the proportion of genes with a genuine effect^63^. Estimates were obtained with the natural cubic spline method of Storey and Tibshirani^63^, which extrapolates the tuning parameter λ toward 1 to minimise bias, and, as a check, with the bootstrap λ-selection procedure of Storey, Taylor and Siegmund^64^, which instead selects λ to minimise estimated mean squared error; the two gave closely similar values. π₀ was estimated identically for the hemocyte Sex:Time interaction and the carcass Sex:Infected interaction, so that the two tissues could be compared directly on datasets of similar size and replication. The asymptotic properties of these estimators hold under weak dependence among tests, a condition that gene expression data are expected to satisfy because genes correlate within pathway-sized blocks that are largely independent of one another^64^.

#### Tissue-specificity of the infection response

To compare the infection response between hemocytes and carcass, adult samples from both tissues were analysed together, restricted to unchallenged flies and the 4 h timepoint so that the two tissues were matched in time, leaving 11,416 genes. Counts were modelled with the design ∼ Sex + Tissue + Infected + Tissue:Infected. Collection date could not be fitted in this model, because two collection days contributed carcass samples only and no unchallenged samples, making date collinear with both tissue and infection status. Sexes were pooled with sex retained as an additive covariate, justified by the absence of a detectable sex-dependent hemocyte response reported above. The infection effect within each tissue was obtained as a single coefficient by fitting the model twice, with carcass and then hemocyte as the reference tissue, so that the two estimates were shrunken on an equivalent footing and could be compared directly on the same axes. Genes were classified by effect size rather than by significance: hemocyte– or carcass-specific if the response exceeded 1 log₂ unit in one tissue while remaining below 0.5 log₂ units in the other, and shared if the two responses differed by less than 0.5 log₂ units with at least one exceeding 0.5.

#### Constitutive sex differences in hemocytes

To identify sex differences that persist independently of infection, hemocyte counts were refitted with an additive design (∼ Date + Sex + Time) in which the sexes are assumed to follow parallel trajectories, so that the Sex coefficient estimates a constant offset between them across the time course. The same filtered counts, metadata and linear time coding were used as for the interaction model above. Genes were called sustained sex-biased at FDR < 0.05 with |log₂ fold-change| > 1, and only where the sex × time interaction from the model above was not significant, so that a single additive offset is a valid summary of the difference; on this criterion one gene was excluded. As a check that the offset is genuinely constant rather than an artefact of borrowing information across timepoints, the sustained estimates were compared with the sex difference at baseline alone, taken from the intercept-level Sex coefficient of the interaction model; the two agreed closely (Pearson r = 0.98).

For display, the volcano p-value axis was square-root transformed and p-values below 10⁻¹⁰⁰ were clamped to that value. Functional categories shown were assigned manually from a curated gene list.

#### Visualisation of individual genes

Normalised counts for individual genes were extracted with *plotCounts* (DESeq2), which divides raw counts by the sample-specific size factors estimated during model fitting. These panels show the data underlying the model estimates, so that induction can be assessed against baseline expression and sample-to-sample variability rather than from fold-changes alone. Counts are plotted on a log₁₀ scale with an independent y-axis range per gene, and lines connect group means.

## Acknowledgements

We wish to thank Steve Jenkins, and the Vale, Obbard, and Agrawal labs for fruitful discussions around this work. We thank Adam Bajgar and Gabriela Krejčová for advice on flow cytometry of adult hemocytes, and the School of Biological Sciences Flow Cytometry Facility at the University of Edinburgh, in particular, Martin Waterfall, Marie Goepp, and Chris Hall for their assistance.

## Supplementary Figures

**Supplementary figure 1:**
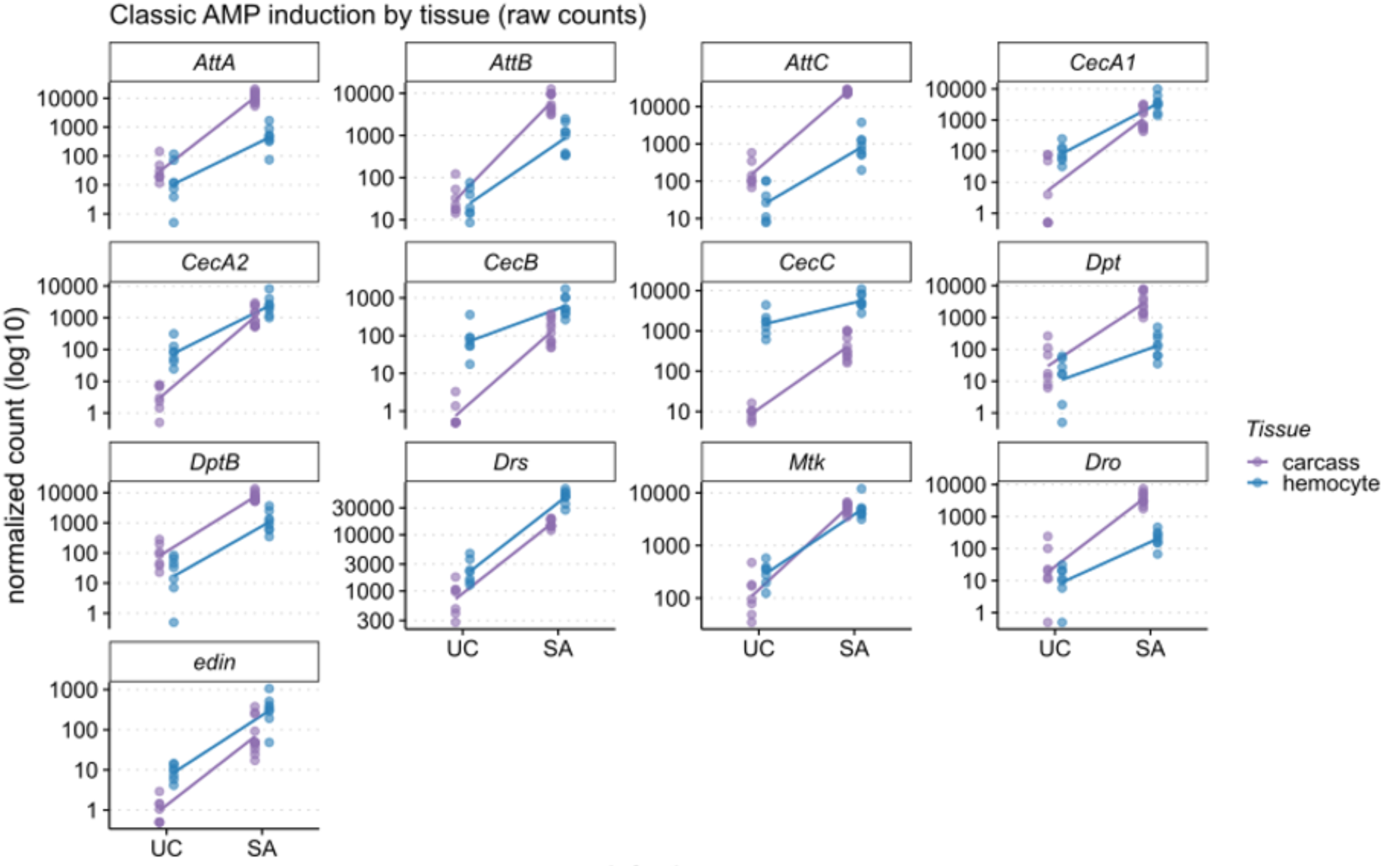
Response of AMP to *S. aureus* injection in carcass vs hemocytes. Normalised counts for the Bomanin in unchallenged (UC) and infected hemocytes at 4 h. Points are individual samples and lines connect group means; y-axis log₁₀ with a free range per gene.

**Supplementary figure 2:**
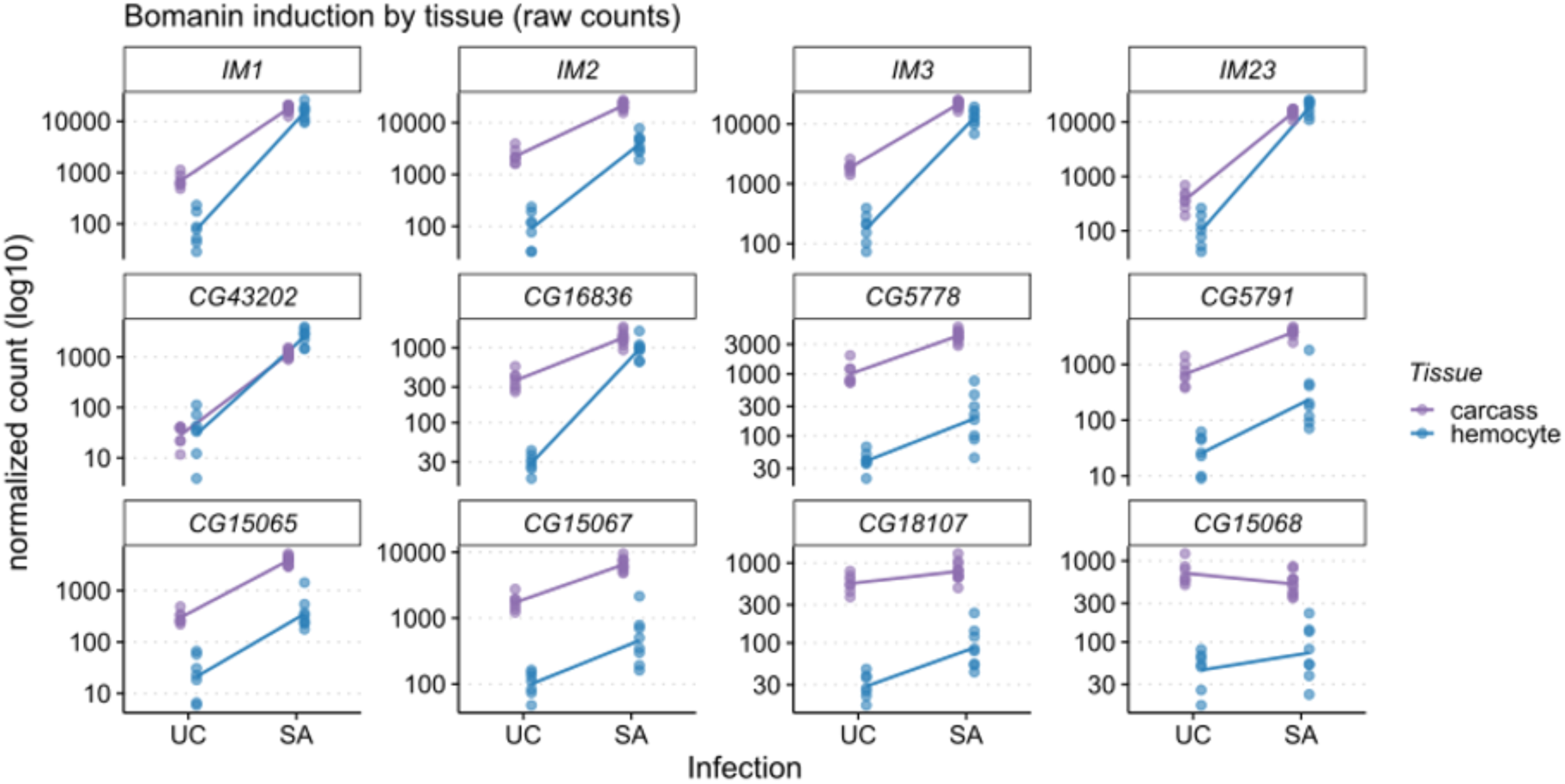
Response of Bomanin to S. aureus injection in carcass vs hemocytes. Normalised counts for the Bomanin in unchallenged (UC) and infected hemocytes at 4 h. Points are individual samples and lines connect group means; y-axis log₁₀ with a free range per gene.

